# Use of a specific set of learner-centered evidence-based teaching practices correlates with higher exam performance across seven STEM departments

**DOI:** 10.1101/2025.06.13.659643

**Authors:** Mallory A. Jackson, Hongjiao Liu, Sungmin Moon, Jennifer H. Doherty, Mary Pat Wenderoth

**Author notes:** These authors contributed equally to this work (Mallory A. Jackson, Hongjiao Liu).

## Abstract

Myriad studies support the claim that active learning improves student academic performance in STEM, yet lecture remains the dominant form of instruction. Faculty offer multiple reasons for not using active learning with many expressing confusion as to what active learning is. Contributing to that confusion is the fact there are a multitude of named active learning methods and varied implementations of each. In an effort to better understand how specific elements of active learning might contribute to enhanced academic performance, we used a more fine-grained classroom observation tool, PORTAAL, to observe teaching practices across 157 STEM courses. We used a principal component analysis to reduce the observed teaching practices to specific sets of practices. We found a continuum of implementation of teaching practices that ranged from instruction-centered to learner-centered. The instruction-centered practices included high Bloom’s level questions, students working alone, instructors answering and explaining questions, and providing alternative answers. The learner-centered practices were volunteer or randomly-called students explaining answers, instructors prompting students’ logic, and instructors giving positive feedback. Using a linear regression to analyze the data from all courses, we found medium and medium-high levels of learner-centered teaching practices correlated with higher student exam performance compared to instruction-centered practices. We observed a similar pattern when analyzing data only from introductory courses. We also analyzed interactions of both binary gender and first generation status with learner-centered teaching on exam scores.

We propose that when instructors use the learner-centered practices, they shift the responsibility of the intellectual work of the problem–the learning–to the students, which may result in improved exam performance. Even moderate levels of these learner-centered practices have a positive correlation with exam performance. Faculty can incorporate these key elements of learner-centered practices into their own teaching based on their comfort level, course content, and abilities of their students.

## Introduction

There are myriad studies that support the claim that active learning improves student academic performance in STEM [1–9] and non-STEM courses [10–14]. Though there is strong evidence to support the benefits of implementing active learning in the STEM classroom, lecture remains the dominant form of instruction in STEM [15–17]. Faculty provide many reasons for why they have not implemented active learning in their classrooms. Some state that these new teaching methods consume too much class time at the expense of covering course content [18]. Others cite a lack of incentive to invest the time needed to redesign their courses and to learn a new teaching method. Finally, there is a large cohort who express confusion as to what active learning is and what it looks like in practice [19–23].

“What is active learning?” is, in fact, an important question. Recent articles by Driessen et al., [24] and Lombardi et al., [25] acknowledge that active learning is often poorly defined by education researchers and can encompass a wide variety of teaching methods. Two of the most highly cited meta-analyses on the impact of active learning on student academic performance acknowledge that active learning was often minimally described in the papers in their study [1,2]. Studies were coded as active learning if they meet the following criterion: “Active learning engages students in the process of learning through activities and/or discussion in class, as opposed to passively listening to an expert. It emphasizes higher-order thinking and often involves group work” [1]. Given these broad criteria and that many of the included studies offered vague descriptions of the type of teaching in the study, the actual type of active learning used in each study was not analyzed. A follow up meta-analysis of 41 studies published between 2010 and 2016 found that a high intensity of active learning (use of active learning > 66% of class time) improved student exam performance more than medium or low levels of active learning [2]. This second study collected and analyzed different types of named active learning methods such as flipped classes, in-class group work, and worksheets. However, the authors urged caution in interpreting results as the active learning types reported in the studies were “author-defined and represent general characteristics’’ [2].

In their 1991 book, *Active Learning: Creating Excitement in the Classroom*, Bonwell and Eison defined active learning as “a method of learning in which students are actively or experientially involved in the learning process” [26]. The philosophical foundation of active learning is socio-cultural constructivism which posits that learners acquire new knowledge by working with others to construct new understanding through social discourse and integrating that understanding into their existing bodies of knowledge [27,28]. Basic characteristics of active learning that Bonwell and Eison noted are that students need to be involved in doing things and thinking about what they are doing. That “doing” can be reading, writing, discussing or solving problems and students need to be engaged in the challenging tasks of analysis, synthesis and evaluation [26]. Since Bonwell and Eison’s seminal work in 1991, a multitude of named active learning methods have been created. These methods include but are not limited to Cooperative Learning [29], Collaborative Learning [30], Team Based Learning [31], Problem Based Learning [32,33], Peer Instruction [34], Jigsaws [29], think-pair-share [35], in-class clicker questions [36], Process Oriented Guided Inquiry Learning (POGIL)[37], high structure classes [38], and flipped classrooms [39].

As researchers seldom fully describe the active learning methods they are assessing in their research, it is challenging to determine if one method of active learning is more effective at enhancing student learning than another. As a result of this lack of clarity, there have been calls to begin second-generation research on active learning practices in undergraduate classrooms [1,40]. This new area of research would not only identify key elements of active learning teaching practices, but also determine the dosage of those elements that correlate with increased student exam performance across demographic groups of students.

In an effort to better understand how certain elements of active learning might contribute to enhanced student academic performance, we created and validated the Practical Observation Rubric To Assess Active Learning (PORTAAL)[41]. To develop PORTAAL, we surveyed the education research literature and identified elements of active learning that when tested individually against more traditional passive learning methods had evidence of significant improvement in student academic performance. PORTAAL identifies the elements of active learning that can be observed in the classroom that are research-supported teaching methods. Therefore, throughout this paper we will use the term “PORTAAL practices” to denote teaching practices that are associated with active learning and that are evidence-based.

Previously, we used PORTAAL to collect data from a large set of introductory biology courses at an R1 institution to determine which individual PORTAAL practices correlated with improved student exam performance [42]. We found that student exam scores correlated with the use of four individual PORTAAL practices: small group work, randomly calling on students for answers, providing alternative answers to questions, and the total time students engaged in class. As the use of these practices increased, so did student exam scores. We were aware that instructors were implementing more than one of the PORTAAL practices during their class sessions. However, given our sample size, we were unable to analyze if instructors preferentially used different sets of PORTAAL practices and if those sets of practices had a differential impact on student academic performance.

To expand on our previous research, we conducted an analysis to determine if we could identify sets of PORTAAL practices used in the 146 upper- and lower-division courses in seven STEM departments. We reasoned, if we could identify sets of PORTAAL practices, we could use the sets to investigate how these sets correlate with student exam performance. The research questions for this study are as follows:

RQ1. Can we identify sets of correlated PORTAAL practices based on their use in courses across seven STEM departments?
  RQ1a: Does the use of PORTAAL practice sets vary by STEM department?
RQ2. Does the use of these sets of PORTAAL practices correlate with student exam performance?
  RQ2a. Do the correlations vary by student demographic groups?

## Methods

### Data collection

#### Classroom observation data

Human subjects research was conducted with the approval of the Institutional Review Board at the institution where the study took place (STUDY00002830). Participants were recruited between July 2017 and March 2020. Faculty participation in the study was voluntary and documentation of consent was waived for faculty. Consent was waived for students. Students had an opt-out procedure as described in the IRB protocol if they did not want their data included in the study.

A subset of faculty from seven STEM departments (biology, chemistry, computer science and engineering [CSE], mathematics, physics, psychology, and public health) volunteered to have their courses observed as a part of this study. A total of 157 lower- and upper-division courses, spanning from Winter Quarter 2017 to Winter Quarter 2020, were included in the study. Classroom observations of the 157 courses were conducted in the same manner as in Jackson et al. [43]. Video recordings of class sessions were collected using lecture capture technology. On average, four videos were randomly selected to be coded from each course, totaling approximately 600 individual video recordings. We randomly selected the videos by dividing the ten-week quarter into four equal time periods excluding the first and last week of the quarter and randomly selecting one video from each time period. Pairs of trained researchers coded classroom videos using the PORTAAL classroom observation tool [41]. The PORTAAL coding rubric can be found in the supporting information (S1 Appendix). Two coders individually coded the entire video, then met to discuss any disagreements in coding and came to complete consensus on all codes. After coding, we calculated the average value for each of the PORTAAL practices in each course by totaling the number or duration of each practice and dividing by the number of videos coded for that course. To standardize comparisons, these averages were adjusted to a 50-minute class session, aligning with the institution’s standard class length.

Given the presence of over 30 individual PORTAAL practices in the standardized dataset, we aggregated closely-related practices and removed those that were infrequently observed to ensure the power and interpretability of the analysis (S3 Table). First, we aggregated some of the practices as they measured similar classroom behaviors. For example, students’ volunteered responses (“Vol_Ans_Exp”) represents the combined number of times two similar events occurred in class: students volunteered answers or students volunteered an explanation for their answers (see S3 Table for further explanation of PORTAAL practices). Second, we removed variables that were rarely observed in the sample of 157 courses. For example, instructors providing negative feedback to students was seldom observed across all of the courses, so it was not included in the final set of practices. The final set of ten PORTAAL practices used for analysis is found in Table 1.

**Table 1.** PORTAAL practices used for analysis.

| **PORTAAL Code** | **Long Description** | **Combined Variable** |
| --- | --- | --- |
| HB | Number of high Bloom’s level activities | No |
| Alone | Number of times students worked individually during activities | No |
| SG | Number of times students worked in small groups during activities | No |
| Ins_Ans | Number of times the instructor gave answers during the debriefs | No |
| Ins_Exp | Number of times the instructor explained answers during the debriefs | No |
| Vol_Ans_Exp | Number of times student volunteers gave answers and/or explained answers during the debriefs | Yes |
| RC_Ans_Exp | Number of times randomly called students gave answers and/or explained answers during the debriefs | Yes |
| Prom_Log | Number of times the instructor prompted the students to use logic when thinking about or answering a question | No |
| Alt_Ans | Number of alternative answers that the instructor or students explained | No |
| Pos_FB | Number of times the instructor gave positive feedback | Yes |

#### Exam data

We attempted to collect student exam scores from each of the courses in the study, but some instructors did not provide exam data or did not administer exams in their courses. As a result, our final data set only included the 146 courses for which we had both classroom observation data and exam data. Among the included courses, the number of exams per academic quarter ranged from one to ten, though the majority of courses had three to five exams per quarter. Prior to the analysis, we calculated an average standardized exam score for each student per course by totaling the number of points each student earned on all exams in the course, dividing by the total possible exam points in the course, and multiplying by 100 to standardize the score. Boxplots of standardized exam score distributions for each STEM department are shown in Fig 1.

**Fig 1.**
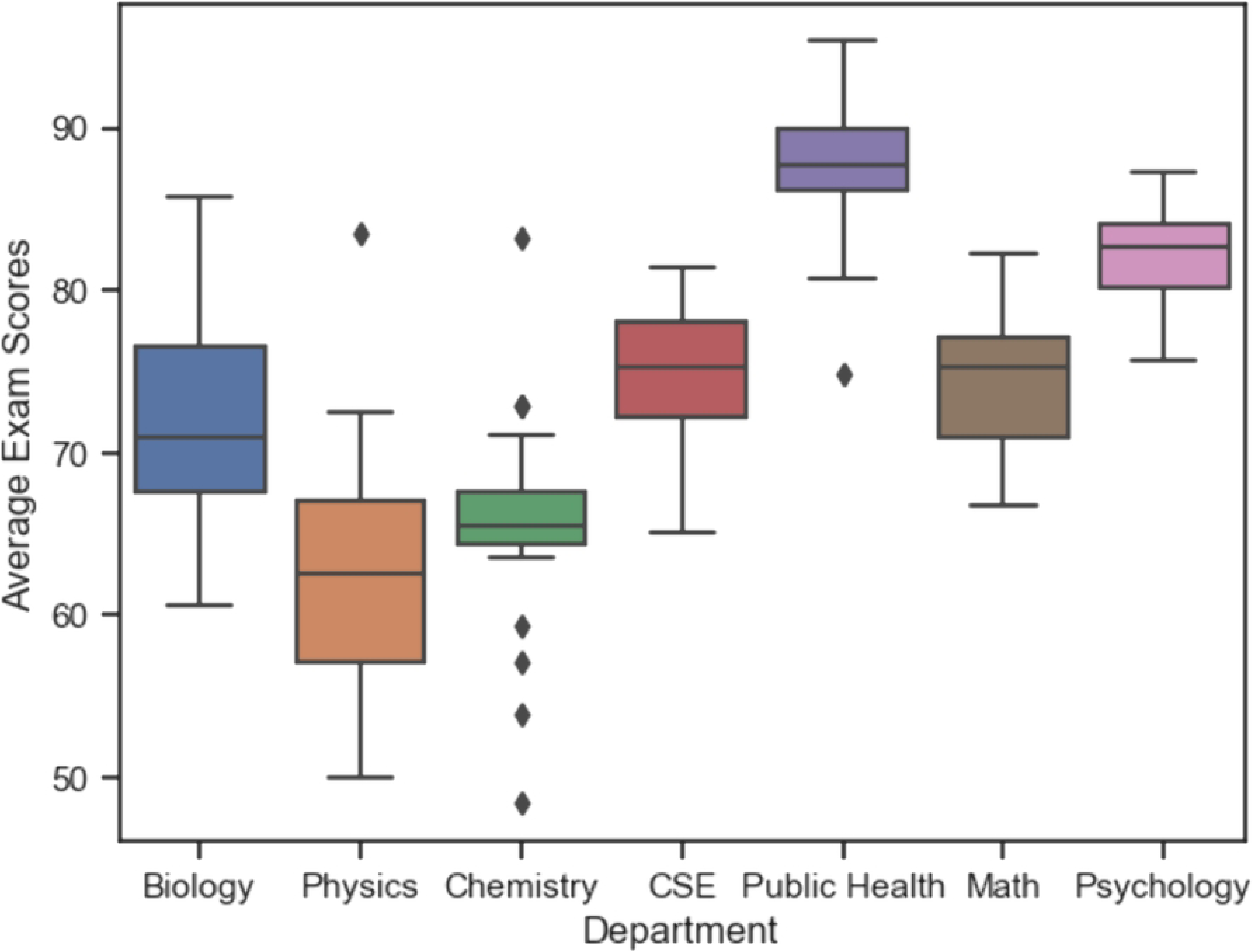
Average exam scores for courses in each STEM department.

Some students enrolled in multiple courses, and therefore had multiple course records and associated PORTAAL values in our dataset. We will refer to each instance of student data and the associated PORTAAL values from that course as a *student data point*. Student data points with any missing exam scores were removed prior to analysis. There were a total of 27,114 student data points in the 146 courses included in the analysis. S4 Table shows the number of student data points, the total number of courses and introductory courses among them, and the number of instructors for each department. These student data points represented 15,990 unique students; 43% of students took more than one course. Lower-division courses ranged from 200 to 700 students per course, whereas upper-division courses ranged from 24 to 120 students. The number and percentage of upper- and lower-division courses can be found in the supporting information (S5 Table).

#### Student demographic information

Student demographic information and GPA at the start of the quarter were obtained from the university registrar. Demographic factors included self-identified binary gender, if the student was the first in their family to graduate from college (first generation status or FGN), if their race/ethnicity is underrepresented in STEM (underrepresented minority or URM: e.g., Hispanic/Latino, Black/African-American, Pacific Islander, Native Hawaiian, Native American), and if they qualified for the Educational Opportunity Program for students identified by the university as economically or educationally disadvantaged (EOP). Descriptive statistics of students in each group are reported in the supporting information (S6 Table).

### Data Analysis

#### Principal Component Analysis

Prior to conducting a principal component analysis (PCA), we first computed a pairwise correlation matrix to assess collinearity among PORTAAL practices (S7 Fig). Although most practices were positively correlated, no correlation exceeded 0.8, suggesting no severe multicollinearity. Consequently, all selected PORTAAL practices were included in the PCA.

To identify potential underlying sets of correlated PORTAAL practices, we conducted principal component analysis (PCA) as an exploratory analysis. PCA is a dimension reduction approach that summarizes the variation in the original PORTAAL practices using a small number of new variables called principal components (PCs). Each PC is a linear combination of the original variables, and the PCs are uncorrelated with each other. The first few PCs capture the majority of variation within the original data. In the context of our analysis, these PCs could be interpreted as teaching patterns that are characterized by a set of PORTAAL practices.

We performed the PCA on the PORTAAL practices data at the course level. Each data point in the PCA analysis corresponded to a distinct course, with 146 courses in total. We examined the contributions of the PORTAAL practices to the top PCs by inspecting the loading matrix, and further examined how the courses cluster by department based on the PC values. We used a variance cutoff of 60% to decide the number of PCs to retain in our analysis. Based on this cutoff value, we used the first two PCs (PC1 and PC2) from the PCA analysis of PORTAAL practices as variables in subsequent regression models for exam scores.

PC1 and PC2 are a continuum of values representing variation in the number of times PORTAAL practices were used in the courses. As we were interested in how broad, categorical levels that could represent low, medium, medium-high, and high use of the sets of teaching practices correlated with exam scores, we binned the observed values of each PC into quartiles to be used in the subsequent analyses. For both PC1 and PC2, quartile 1 (Q1) represents the lowest values, quartile 2 (Q2) represents medium values, quartile 3 (Q3) represents medium-high values, and quartile 4 (Q4) represents the highest values.

We conducted the PCA analysis using the *stats* package in R (version 4.0.2).

#### Regression Modeling

We performed regression modeling using linear mixed models (LMM) to determine how PC1 and PC2 correlated with student exam scores. We conducted the analyses with LMM models in R (version 4.0.2) using the *lme4* and *lmerTest* packages. The PC1 and PC2 quartiles (*tp_PC1_quartile* and *tp_PC2_quartile* in the formula below) were included as predictors of interest and treated as fixed effects, and the average exam score (*Average*) was the outcome. We included four intercept random effects at different clustering levels to account for the semi-nested structure of the data, where student exam scores within the same cluster could be correlated with each other (Fig 2). First, to account for the correlation among exam scores from students taught by the same instructor, we included the instructor ID (*Instr_ID*) as a random effect. Secondly, to account for the correlation among exam scores from the same course, we included the course code (*Course_Short*) as a random effect. Furthermore, the same course could be offered in multiple academic quarters/terms, so we included a random effect for the unique offering of a course during a quarter (*Course_Short_ Quarter*) to account for the correlation among exam scores from the same course in the same term. Finally, to account for the correlation among exam scores from the same student, we included the student ID (*SID*) as a random effect.

**Fig 2.**
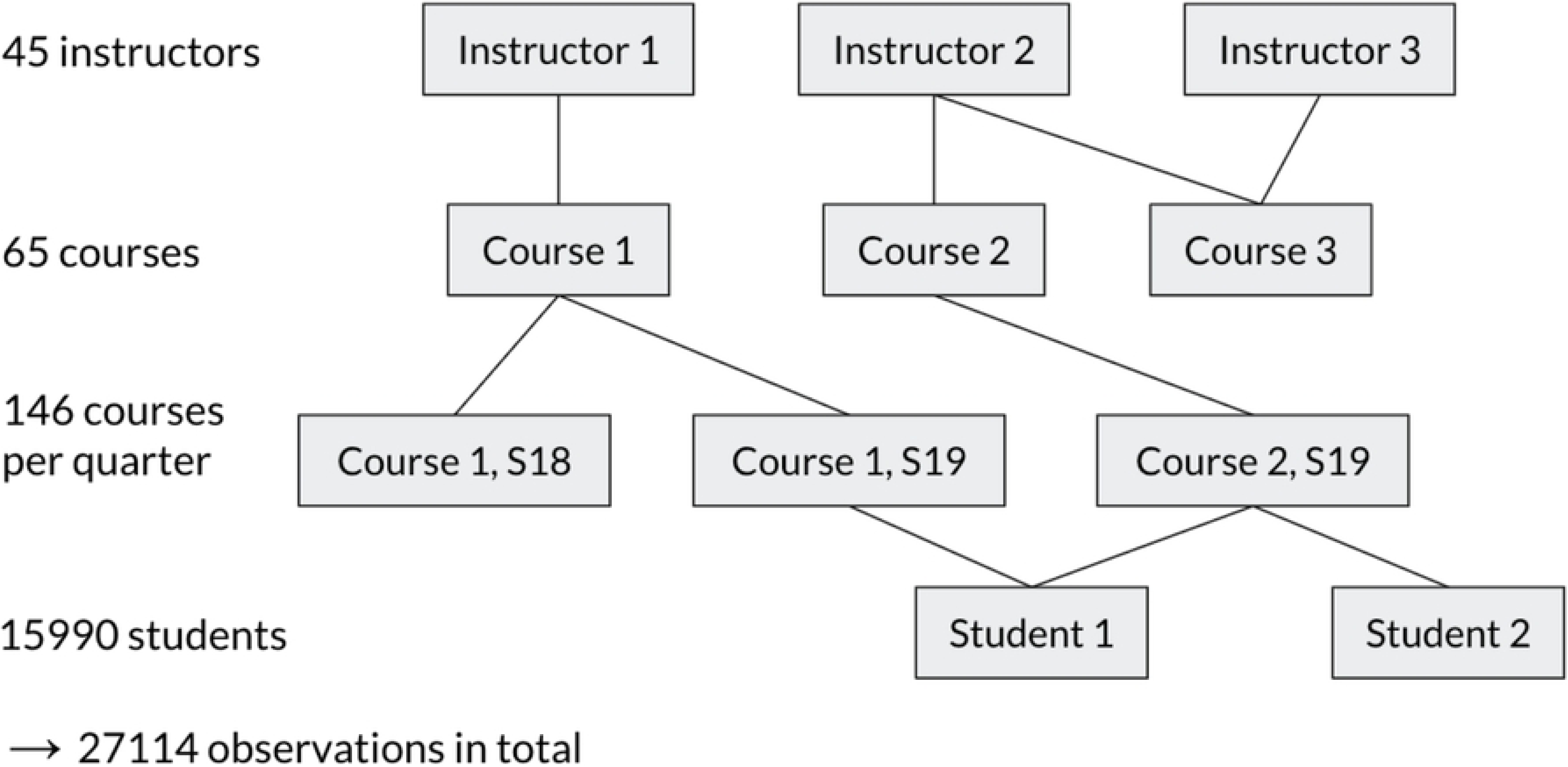
Semi-nested structure of the data.

In several cases, a single instructor taught more than one course, and some courses were taught by multiple instructors, either within the same term or in different terms. Additionally, students could be enrolled in more than one course. As a result, the data had a semi-nested structure rather than a strictly nested one. We included the following covariates as fixed effects in the regression model: students’ incoming GPA (*GPA_Start*) for each course, gender (*Gender*), EOP status (*EOP*), URM status (*URM*), first generation status (*FGN*), and course level (upper- or lower-division; *Course_Level*). As different departments could adopt systematically different teaching approaches and have distinct exam difficulties, the department was likely a confounding factor in the association between PORTAAL practices and exam scores. We also observed a large variation in average exam scores by department as shown in Fig 1. Therefore, we also included the department (*Department*) as a fixed effect in the regression model. The full regression model for the primary analysis is as below:

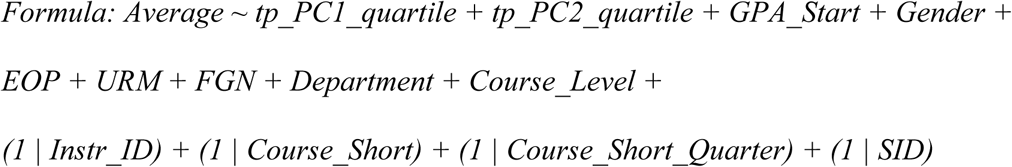

The variance explained by fixed effects was calculated according to Nakagawa and Shielzeth’s method [44], and the variances explained by random effects and the residual variance were obtained from the variance component estimates of the LMMs.

We conducted a subgroup regression analysis for introductory courses to determine if the correlation of PC1 and PC2 quartiles with exam scores was different for these courses than in the overall analysis. Courses were coded as introductory if they were lower-division courses required to complete STEM majors (e.g., introductory biology, introductory chemistry, algebra-based physics, introductory computer programming). For this analysis, the regression model remained the same as above but the data set was reduced to introductory courses only.

We conducted additional analyses to examine the interaction between demographic variables of interest (binary gender and FGN status) and PC1 and PC2 quartiles on exam scores. The general formulas for the regression models with interaction effects are as below:

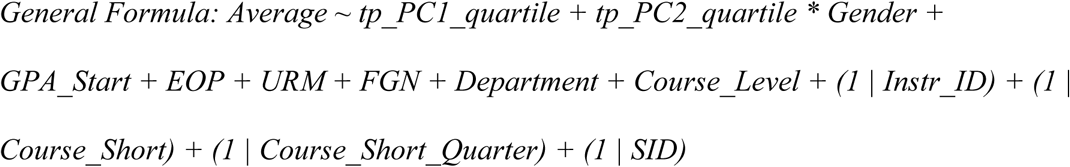

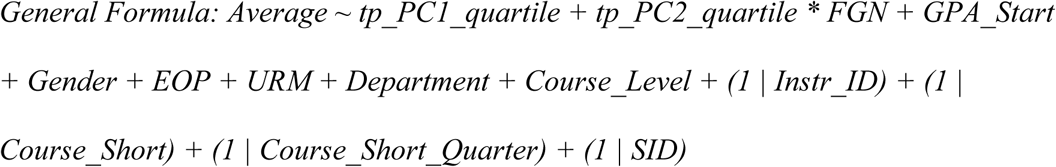

## Results

### RQ1. Can we identify sets of correlated PORTAAL practices based on their use in courses across seven STEM departments?

A scree plot of the percent of variance explained by each PC showed that the first two PCs explained 61.2% of the total variance in the use of the 10 PORTAAL practices (S8 Fig). In reviewing the results of PCA, it is important to note that only the relative PC magnitudes and relative sign patterns of the loadings, rather than the absolute signs, are meaningful. Positive and negative signs are only meant to represent opposite directions of loading and do not imply that one set of factors is better than or worse than another–they are merely different from each other, not unlike the positive and negative poles on a battery.

Based on the heatmap of the loading matrix (Fig 3), PC1 was positively correlated with all practices, meaning we can interpret PC1 as positively correlated with a teaching pattern that incorporates all 10 practices at varying levels of use. PC2 correlated positively with one group of practices and correlated negatively with another group of practices. The group that was positively correlated with PC2 consisted of the following four practices: volunteers giving/explaining answers, randomly called students giving/explaining answers, the instructor prompting logic, and the instructor giving positive feedback. The group that was negatively correlated with PC2 consisted of five practices: instructors explaining, instructors answering, students working alone, students or the instructor explaining alternative answers, and use of high Bloom’s level in-class questions. The positively correlated PORTAAL practices (red in the heatmap of PC2) represents a more learner-centered teaching pattern, as students rather than the instructor were more actively engaged in the classroom learning environment. The negatively correlated practices (blue in the heatmap of PC2) represents a more instruction-centered teaching pattern.

**Fig 3.**
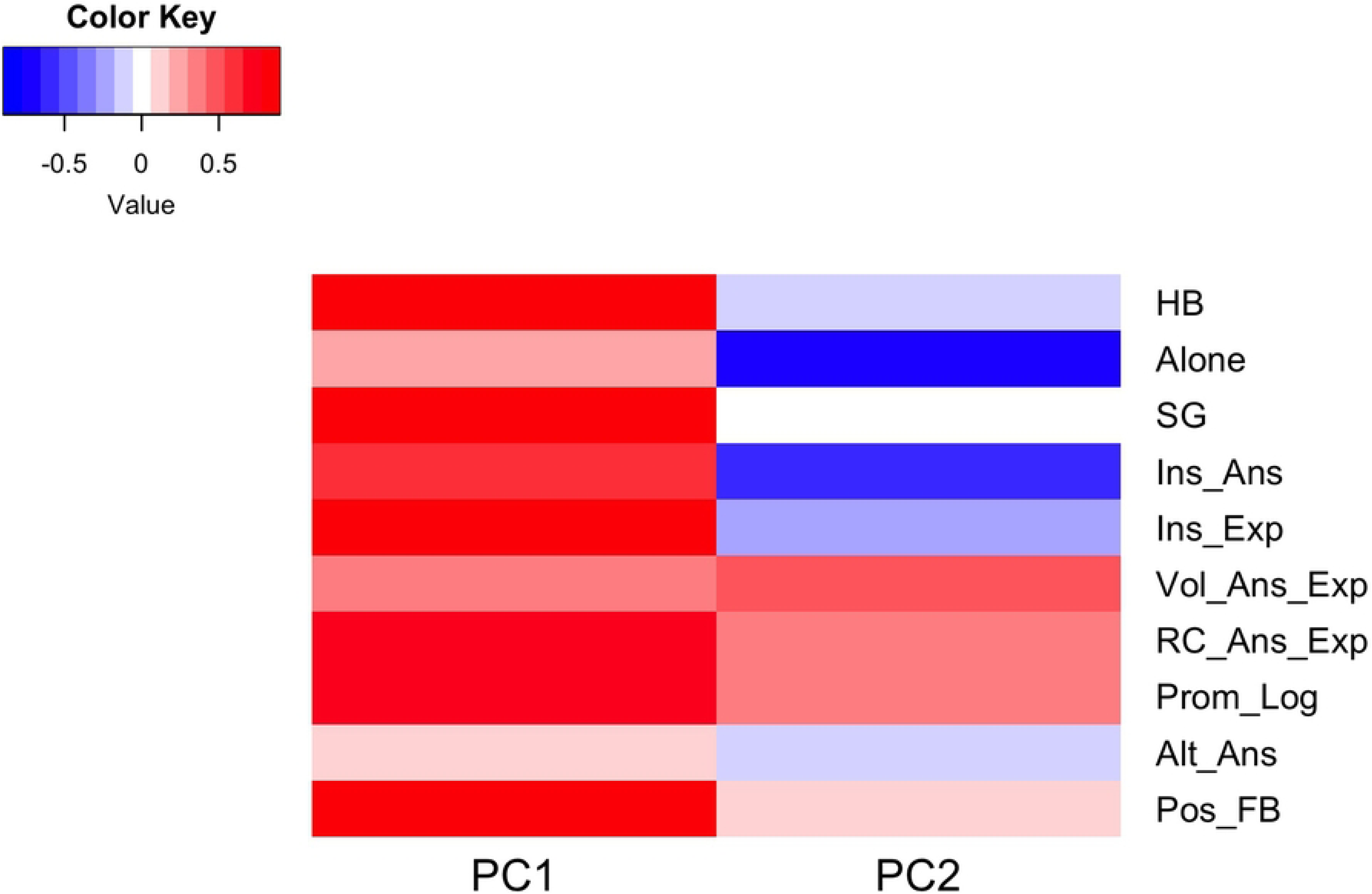
Heatmap of the loading matrix. Each entry represents how much a particular PORTAAL practice contributes to, or correlates with, a particular PC, where darker colors indicate larger contributions. Red represents a positive correlation and blue represents a negative correlation.

### RQ1a: Does the use of PORTAAL practice sets vary by STEM department?

To visualize how the sets of PORTAAL practices were used in each course in each department, we plotted the PC1 values against the PC2 values, colored by department (Fig 4). The PC1 values on the x-axis represent a positive linear combination of the use of all 10 PORTAAL practices. A higher PC1 value is associated with a higher overall use of the 10 practices. The PC2 values on the y-axis represent the variation in use of learner-centered versus instruction-centered sets of PORTAAL practices. Higher PC2 values represent higher use of the set of four more learner-centered practices. Smaller PC2 values represent higher use of the set of five more instruction-centered practices. For example, a course with a PC2 value of -3 used more instruction-centered practices on average than a course with a PC2 value of -1.

**Fig 4.**
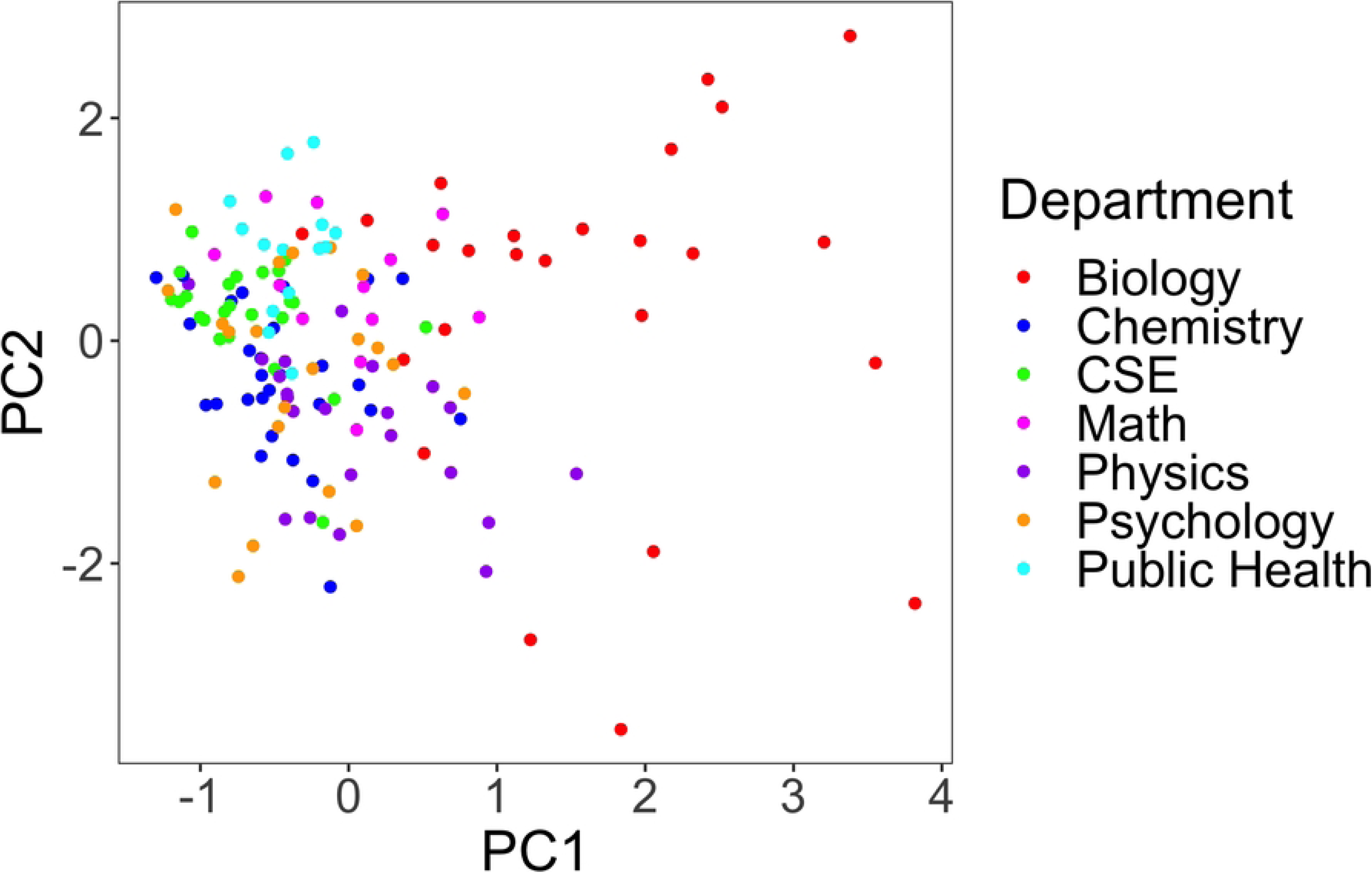
Scatter plot of PC1 and PC2 values for each course, color-coded by department. PC1 (x-axis) represents the overall use of all 10 PORTAAL practices, from low use (-2) to high use (4). PC2 (y-axis) represents the use of instruction-centered and learner-centered sets of practices, with higher use of instruction-centered practices less than zero, and higher use of learner-centered practices greater than zero.

Courses tended to cluster by department, indicating that there are similar teaching styles within departments and different patterns of teaching across departments. Courses in the biology department seem to have a vastly different teaching pattern compared to other departments: they adopt more teaching practices overall (based on the larger PC1 values on the x-axis compared to other departments) and have large variations in learner-centered versus instruction-centered teaching styles based on the wide range of PC2 values on the y-axis. Courses in psychology had lower average PC1 values indicating less overall use of PORTAAL practices, but also had a large range in PC2 values. Physics courses had more variation in PC1 values, which indicates a range of total use of PORTAAL practices, but most of these practices were instruction-centered, as indicated by the lower PC2 values. CSE, public health, and chemistry courses had lower average use of all PORTAAL practices, and CSE and public health courses used more learner-centered practices on average than in chemistry.

### RQ2. Does the use of these sets of PORTAAL practices correlate with student exam performance?

We conducted a linear regression analysis using data from all the courses to determine how PC1 and PC2 correlated with student exam scores. Next, we conducted a subgroup analysis using only the data from the introductory courses. We conducted follow-up linear regression analyses to determine if there was an interaction between demographic variables, binary gender and first generation status of the students, and PC1 and PC2 quartiles on exam scores.

#### Analysis of All Courses

We fitted the LMM model as described in the methods to the full data set with all courses to determine the correlation between PC1 and PC2 quartiles and exam scores. Table 2 shows the percent variance explained by the fixed effects and each of the random effects, out of the total variance in student exam score. The fixed effects, including the teaching practices and covariates, accounted for 47.3% of the total variation in student score. Among the random effects, variation across students accounted for the most variation (11.1%), followed by variation across instructors, courses, and unique course offerings (Table 2). Even after including fixed effects and random effects, there was still 25.4% residual variation in student exam scores that was not accounted for.

**Table 2.** Percent variance in student exam scores explained by the fixed and random effects in the LMM from the primary analysis.

| Component | Fixed effects | Student | Instructor | Course | Course Quarter Pair | Residual |
| --- | --- | --- | --- | --- | --- | --- |
| Variance explained | 47.3% | 11.1% | 7.8% | 5.9% | 2.5% | 25.4% |

The results of the LMM for the correlation between PC1 or PC2 quartiles and student exam scores from all the observed courses are reported in Table 3. Quartile 1, representing the lowest PC values for each PC, was used as the reference group in the analysis. The overall results indicate no significant correlation between the PC1 quartiles and student exam scores when controlling for the PC2 quartiles, covariates, and random effects. This suggests that the overall usage of the 10 PORTAAL practices at any level or intensity represented by the four quartiles do not contribute significantly to the variation in student exam scores across all STEM departments.

**Table 3.** Results from the overall LMM for PC quartiles.

| Principal Component | Quartile (Comparison: Q1) | Student Data Points | Estimate | 95% Confidence Interval | *p*-value |
| --- | --- | --- | --- | --- | --- |
| PC1 | Q1 | 7,915 | - | - | - |
|  | Q2 | 6,555 | 0.22 | [-1.50, 1.94] | 0.8014 |
|  | Q3 | 6,022 | -0.42 | [-2.39, 1.55] | 0.6786 |
|  | Q4 | 6,622 | 1.21 | [-1.88, 4.31] | 0.4438 |
| PC2 | Q1 | 9,054 | - | - | - |
|  | Q2 | 6,608 | 3.08 | [1.48, 4.69] | 0.0002*** |
|  | Q3 | 5,910 | 3.85 | [1.48, 6.23] | 0.0019** |
|  | Q4 | 5,542 | 1.29 | [-1.63, 4.21] | 0.3897 |

*Note.* \* $p < 0.05$ , \*\* $p < 0.01$ , \*\*\* $p < 0.001$ .
*Covariates in the model included students' incoming GPA for each course, gender, EOP status, URM status, first generation status, and course level.*

However, the PC2 quartiles were correlated with student exam scores. Compared to a low level of learner-centered and a high level of instruction-centered practices (Q1 of PC2), a medium level of learner-centered practices (Q2 of PC2) is associated with an increase of 3.08 exam points (*p* = 0.0002) on average. The same holds for a medium-high level of learner-centered practices (Q3 of PC2), with an increase of 3.85 exam points (*p* = 0.0019) on average compared to a low level of learner-centered and a high level of instruction-centered practices. The highest intensity of learner-centered and lowest level of instruction-centered practices, represented by quartile 4 of PC2, was not significantly correlated with exam scores.

#### Analysis of Introductory Courses

We conducted a subgroup regression analysis to determine how the PC values correlated with exam scores in introductory STEM courses. The results of the LMM for introductory courses are reported in the supporting information (S9 Table). We found that, in general, the results followed the pattern of the overall results where a medium level of learner-centered activity (Q2 of PC2) was correlated with higher exam scores (3.24 points; *p* = 0.0003). However, the association between a medium-high level of learner-centered activity (Q3 of PC2) and exam scores was not significant (2.19 points; *p* = 0.0899), unlike in the overall analysis.

### RQ2a. Do the correlations vary by student demographic groups?

We conducted follow-up regression analyses to determine if the association between teaching patterns, as measured by PC1 and PC2 quartiles, and exam scores was different for female students compared to male students, or for students who were the first in their family to graduate from college compared to non-first generation students.

#### Interaction Analysis for Gender

There was no significant interaction between gender and PC1 quartiles in terms of association with student exam scores. In other words, there was no evidence that the overall usage of 10 PORTAAL practices at varying intensities (quartiles) was associated with differential exam performance for female and male students. There was a significant interaction between gender and the quartiles of learner-centered vs. instruction-centered activity (PC2) on student exam scores (*p* < 0.001 from likelihood ratio test). Table 4 shows the results of the interaction of PC2 and gender on student exam scores. In particular, the average increase in exam score when comparing a medium level of learner-centered activity (Q2 of PC2) to a low level of learner-centered activity (Q1 of PC2) was significantly larger in male students than in female students (3.78 versus 2.59 exam points; *p* = 0.0001). The difference between male students and female students’ exam scores was not significant when comparing a medium-high or high level of learner-centered activity (Q3 or Q4 of PC2) to a low level of learner-centered activity (Q1 of PC2).

**Table 4.** Results from LMM with gender by PC2 quartile interaction on student exam scores.

| Gender | Quartile<br>(Comparison: Q1) | Estimate | 95% CI | *p*-value<br>(interaction term) |
| --- | --- | --- | --- | --- |
| Female | Q2 | 2.59 | [0.96, 4.22] | - |
|  | Q3 | 3.74 | [1.34, 6.13] | - |
|  | Q4 | 1.44 | [-1.50, 4.37] | - |
| Male | Q2 | 3.78 | [2.13, 5.43] | - |
|  | Q3 | 4.01 | [1.61, 6.41] | - |
|  | Q4 | 1.01 | [-1.94, 3.96] | - |
| Female vs.<br>Male | Q2 | -1.19 | [-1.81, -0.58] | 0.0001*** |
|  | Q3 | -0.27 | [-0.92, 0.38] | 0.4111 |
|  | Q4 | 0.42 | [-0.23, 1.07] | 0.2023 |

*Note.* \* $p < 0.05$ , \*\* $p < 0.01$ , \*\*\* $p < 0.001$ .
*Covariates in the model included students' incoming GPA for each course, EOP status, URM status, first generation status, and course level.*

#### Interaction Analysis for First Generation Status

There was no significant interaction between first generation status and PC1 quartiles in terms of association with student exam scores. Compared to a low level of learner-centered activity (Q1 of PC2), both a medium level of learner-centered activity (Q2 of PC2) and a medium-high level of learner-centered activity (Q3 of PC2) were associated with higher exam scores in both FGN and non-FGN students (Table 5). The overall association between the intensity of learner-centered vs. instruction-centered activity (PC2) and student exam score differed between FGN and non-FGN students (p-value < 0.001 from likelihood ratio test). The average score increase comparing a medium-high level of learner-centered activity (Q3 of PC2) to a low level of learner-centered activity (Q1 of PC2) was significantly larger in non-FGN students than in FGN students (4.03 exam points versus 3.21 exam points; *p* = 0.0234). In contrast, the average score increase comparing a high level of learner-centered activity (Q4 of PC2) to a low level of learner-centered activity (Q1 of PC2) was significantly larger in FGN students than in non-FGN students (1.85 exam points versus 1.03 exam points; *p* = 0.0173), although the score increase by itself was not significantly different from zero in either non-FGN or FGN students.

**Table 5.** Results of LMM with FGN by PC2 quartile interaction on student exam scores.

| FGN Status | Quartile (Comparison: Q1) | Estimate | 95% CI | P-value (interaction term) |
| --- | --- | --- | --- | --- |
| FGN | Q2 | 3.31 | [1.64, 4.99] | - |
|  | Q3 | 3.21 | [0.78, 5.64] | - |
|  | Q4 | 1.85 | [-1.11, 4.80] | - |
| non-FGN | Q2 | 3.00 | [1.38, 4.61] | - |
|  | Q3 | 4.03 | [1.65, 6.40] | - |
|  | Q4 | 1.03 | [-1.90, 3.95] | - |
| FGN vs. non-FGN | Q2 | 0.32 | [-0.33, 0.96] | 0.3334 |
|  | Q3 | -0.81 | [-1.52, -0.11] | 0.0234* |
|  | Q4 | 0.82 | [0.15, 1.50] | 0.0173* |

*Note. \* $p < 0.05$ , \*\* $p < 0.01$ , \*\*\* $p < 0.001$ .*
*Covariates in the model included students' incoming GPA for each course, gender, EOP status, URM status, and course level.*

## Discussion

Ours is one of the first studies to contribute to the new area of second-generation active learning research. This new area of education research calls for not only identifying key elements of active learning teaching practices, but also for determining the dosage of those elements that correlate with increased student exam performance across demographic groups of students [1,40]. Our research has demonstrated a significant correlation between a specific set of PORTAAL teaching practices (i.e., learner-centered practices), the intensity of use of those practices, and increased student exam performance.

One of the major challenges of active learning research is that active learning teaching methods encompass a wide variety and combination of classroom activities, called by many different names and implemented in a variety of ways. The PORTAAL classroom observation tool identifies specific teaching practices that when used alone are documented in the literature to improve student academic performance [41]. The PORTAAL tool therefore provides a more granular analysis of teaching methods and their correlation with student academic performance.

Our previous work quantified how individual PORTAAL practices correlate with student exam performance across 33 biology courses at an R1 institution [42]. Our current work extends this research in three ways; we have determined 1) sets of PORTAAL practices used in courses 2) how the sets and intensity of use of these sets of PORTAAL practices correlate with student exam scores, and 3) we have extended our findings beyond undergraduate biology courses to include six other undergraduate STEM disciplines.

### RQ1. Can we identify sets of PORTAAL practices based on their use in courses across seven STEM departments?

We found that instructors used the 10 PORTAAL practices at varying levels of use while teaching their courses, and some practices were used at a higher intensity than others. This set of 10 PORTAAL practices are relatively common teaching practices used in some combination in many of the named active learning techniques that include but are not limited to think-pair-share [35], peer instruction [45], traditional clicker question format [36,46], and high structure classes [38]. Given the widespread calls to move away from traditional lecture-based teaching and create more interactive STEM classrooms that fully engage students in the learning process [47–50], as well as the growing efforts to train and support faculty in making these changes [51–56], we were pleasantly surprised to observe that many instructors who volunteered to be a part of this study were using at least low levels of these 10 teaching practices. These results may suggest that active learning is becoming a part of the pedagogy of STEM courses at this university.

Our analysis also found two different sets of teaching practices. One set of practices more often included PORTAAL practices we have designated as instruction-centered while the other set included practices that we have designated as learner-centered. Individually, each of the 10 PORTAAL practices in this study has evidence in the literature that the practice supports improved student learning, as compared to traditional lecture-based teaching [41]. However, our results indicate that use of a specific set of these practices correlates with an increase in academic performance as compared to the other set of practices.

To illustrate how these sets of teaching practices appeared in the classrooms we observed, we constructed representative teaching scenarios based on common patterns seen across the classroom videos. These scenarios highlight the typical sequences of instructor and student actions associated with the learner-centered and instruction-centered sets of practices. In the learner-centered classroom, the practices of students volunteering their answers, randomly calling on students for their answers, the instructor prompting logic, and providing positive feedback to students were used more often. Courses that used more of these practices had higher PC2 scores. A possible teaching scenario in this course would be that the instructor poses a question to the class, prompts the students to use logic when solving the problem, allows time for students to arrive at an answer, and then solicits students to answer and explain their logic by either asking for a student to volunteer their answer or using random call to call on a student or group of students. The instructor closes the activity by providing positive feedback to the class or individual students. We see these practices as actively centering the classroom on the learning experience of the students, which is why we have termed this set of teaching practices, learner-centered.

The teaching practices found in the instruction-centered set include the instructor posing high Bloom’s level activities, students working alone, the instructor answering the problem they posed, the instructor providing the explanation for the answer, and instructors or students providing alternative answers to questions. Courses that implemented more of these practices in their classes had lower PC2 scores. A classroom scenario for the instruction-centered practices could be the instructor asking students to answer a multiple-choice question that involves a higher Bloom’s level analysis question. The students are instructed to first work alone and then use an online response system to select the correct answer from multiple choices. After a period of time, the instructor provides the correct answer with an explanation. The instructor would then proceed to explain why each of the incorrect answers is incorrect.

### RQ1a: Does the use of PORTAAL practice sets vary by STEM department?

There was a continuum of use of all 10 PORTAAL practices, and a range of practices from more instruction-centered to more learner-centered across the seven STEM departments. The department that showed the greatest variation in PC values for courses was the biology department. Instructors in biology courses used the PORTAAL practices more often and ranged from instruction-centered to learner-centered, though the majority of the courses were learner-centered. Many of the faculty in this department have been very involved over the past 15 years in faculty development efforts to implement evidence-based teaching practices in their courses. This department has also had an active discipline-based education research group, which has created a community of support for implementing active learning. Andrews et al. [57] documented the positive influence discipline-based education research faculty have on encouraging colleagues within a department to implement more evidence-based teaching practices.

Of the remaining six departments, physics and mathematics showed a broad range of frequency of use of the practices. However, physics courses appeared to be more instruction-centered while mathematics courses tended toward more learner-centered practices. Nationally, physics has a long tradition of implementing peer instruction, as this teaching practice originated with Eric Mazur, a Harvard physics professor [45]. Therefore, we were not surprised to see some physics courses have a high frequency of use of practices. Peer Instruction often involves thinking and answering alone before discussing with a peer, re-submitting an answer, and then having the instructor provide the answer and explanation as well as discuss alternate answers.

Given the method of Peer Instruction, we expected to see many of these courses fall more in the instruction-centered range. As many of the topics in physics are challenging, it is reasonable to assume the in-class questions were coded as high Bloom’s level. The PORTAAL data from the mathematics courses indicated that instructors included in this study often structured their class sessions around complex, multi-step problems. After assigning the problem to the class, students would often work in groups to solve the problem, and would then be called on to offer answers to each step of the problem. Our finding that the mathematics courses were, on average, more learner-centered than the physics courses may suggest that students in mathematics courses offered more answers and explanations to questions, while in physics, instructors more often provided answers and explanations to in-class questions.

The four remaining departments each showed lower levels of implementation of PORTAAL practices in their courses. However, in public health and CSE courses, the practices that were implemented were more learner-centered, while for chemistry and psychology the practices were more instruction-centered. Based on the classroom observations of public health courses, we noted that there is a strong emphasis on group work and projects which could reflect the emphasis on building community that is promoted in this major on this campus. As group work and projects involve considerable student input, they are by their nature very learner-centered activities. However, group work and projects are more time consuming which could explain the lower number of times these practices were observed.

CSE courses also had low implementation of learner-centered practices but they also had the least variation in the type and intensity of learner-centered practices. This tight clustering of learner-centered courses could be due to the discipline-specific pedagogy instructors use. Many of the CSE courses involved live coding as a teaching tool and often called on students to explain the outcomes of the code which made this a learner-centered practice. However, many of these CSE courses are also prerequisites for upper-division courses in this highly competitive major and faculty may feel time pressures to cover content required in subsequent courses. The pressure faculty feel to cover content is a well documented barrier to implementation of evidence-based teaching practices [18,21,23,58–61] and could account for the lower number of learner-centered practices implemented.

Similar time constraints may be felt by faculty in the chemistry department [62]. The year-long introductory chemistry sequences are made up of very content-heavy courses. Instructors often feel time pressures because they must cover a large amount of material in each class, as this content is often foundational for the courses that follow in the series. The pressure to cover content was stated explicitly by faculty that we interviewed after we observed their courses, and is explored in depth in our previous work [63]. As many of the teaching practices (such as having students answer and explain concepts) may require more class time, faculty may be reluctant to use them and may prefer to focus on explaining the answers themselves which shifts the class into a more instruction-centered format.

Though many psychology courses had a lower frequency of use of the PORTAAL practices, they did show a range of instruction to learner-centered practices. Again, it is possible that the content of the course in this major may influence the teaching methods. Some of the psychology courses in this study were for non-majors. It is possible that when faculty are relieved of the pressure to cover content and have an interest in generating curiosity in learners they may shift to more learner-centered practices. However, some of the courses were for majors and the need to cover content could once again be shifting the teaching practices toward instruction-centered.

### RQ2. Does the use of these sets of PORTAAL practices correlate with student exam performance?

To extend our previous research that focused on how individual PORTAAL practices correlate with exam performance, we used PCA as a dimension reduction approach to identify how the intensity of use of sets of PORTAAL practices correlate with student exam performance. Our analysis of all courses did not show a significant correlation between intensity of use of the 10 PORTAAL practices and student exam scores across all STEM departments.

This could suggest that when these 10 PORTAAL practices are used in concert, they do not contribute significantly to the variation seen in students’ exam scores. These results could indicate there are other factors that play a more significant role in explaining students’ exam performance than this particular set of teaching practices. In our original paper describing the development of the PORTAAL tool [41], we acknowledged that there are many factors that contribute to student learning and academic success including but not limited to pre- and post-class assignments, student motivation, and self-efficacy.

It is also possible that in courses that had increased use of these practices, the instructor may also have increased the cognitive challenge of exams. This is what Freeman et al. found when they analyzed weighted Bloom’s scores of exams across six terms of transforming an introductory biology course from a low level of active learning to a much higher level of active learning [38]. As they added more active learning activities in each subsequent term, they found that the cognitive challenge of their exams also increased. Increasing the cognitive challenge level of exams may diminish the statistical impact of active learning on exam scores.

As we consider it unethical to use traditional lecture as a control condition, we used PCA to explore how different components of active learning correlate to student exam scores and the relative effectiveness of the components. When comparing the learner-centered set of PORTAAL practices to the instruction-centered set of practices, we found as the intensity of learner-centered practices increased, so did the student exam scores. Others have also reported a correlation between increased intensity of use of broadly-defined active learning practices and increased academic performance [38,64–66]. In each of these studies, various types of active learning or course structure such as pre-reading quizzes, practice exams, in-class clicker questions and group work, and use of worksheets were added to courses over time to determine impact on student academic performance. Our study differs from the previous research as we examined how the increased use of a specific set of the PORTAAL practices correlates with student exam scores.

Each of the 10 PORTAAL practices when used alone is documented in the literature to support student learning [41]. However, based on our findings, we propose that when faculty use the set of learner-centered practices (students volunteer their answers, randomly call on students for their answers, prompt students to use their logic, and provide positive feedback to students after the students have answered), the instructor is setting up the problem and the format for student interaction and shifts the responsibility of the intellectual work of the problem–the learning–to the students. The instructor is facilitating the students’ opportunity to practice the scientific skill of explaining, using a combination of volunteer and random call to allow a greater variety of student voices to be heard, and is offering timely positive feedback to encourage the continued intellectual work of learning by the students.

We propose that a key aspect of the set of learner-centered PORTAAL practices is their strong alignment with a sociocultural constructivist theory of learning. Sociocultural constructivism posits that learning occurs as learners interact and collaborate to test and build their understanding [27]. A similar emphasis on the interaction of learners in the learning process is a key component of the ICAP Framework [67]. Chi and Wylie hypothesized and provided evidence that as students become more engaged with the learning process and move from passive to active to constructive and finally interactive engagement, their learning increases. By having students or groups of students answer and explain answers to in-class questions, the instructor has the potential to facilitate a whole class interaction with the learning process.

Each of the PORTAAL practices found in the instruction-centered teaching profile (use of high Bloom’s level questions, students working alone, instructor answers and explains the question) is also documented in the literature to support student learning. For example, Jensen et al. found that using high-level Bloom’s questions during formative assessments resulted in students acquiring deep conceptual understanding of the course material and higher grades [68]. Similarly, based on interviews of students who had engaged in peer instruction that involved first working alone, Nielsen et al. found that students valued the alone time as a crucial opportunity to construct their arguments and explanations [69]. This allowed the students to engage in more fruitful discussion with their colleagues during the peer discussion. Smith et al. compared student performance on a second isomorphic in-class concept question after either peer discussion or instructor explanation or a combination of the techniques [70]. They found the combination approach improved average student performance considerably when compared with either technique alone, indicating the value of instructor explanations in addition to group work.

Why then was this grouping of instruction-centered practices not correlated with improved student exam performance as compared to the learner-centered practices? Though using high Bloom’s level questions on in-class assignments has been shown to be beneficial for many students, these questions may cause anxiety for other students and hinder productive thinking during class. Work by Wright et al. found that female students and students with lower socioeconomic status had lower performance on constructed response exams as the Bloom’s level of the exam increased [71]. Wright et al. proposed that these groups of students may have experienced stereotype threat or had a more fixed mindset. Therefore, instructors may need to better scaffold high Bloom’s questions to counteract these influences.

We propose that though Nielsen et al. found students valued working alone [69], this practice also can minimize student interactions. Driessen et al. found that implementation of group work during class increased student performance by 1.00 standard deviations [72] and Weir et al. found student performance on diagnostic tests were highest in courses that added worksheets during in-class group activities [65]. A key aspect of the socio-constructivist framework of learning is an emphasis on the role of social interaction in the development of reasoning in the learning process [27]. Therefore, it is possible that group work may be more beneficial than alone time for student academic performance.

The instruction-centered activity of the instructor answering and explaining answers to in-class questions also may deny students the opportunity to engage in the scientific process of explaining and interpreting results. Having students practice hypothesizing and explaining has been shown to positively predict students’ felt recognition as a scientist, which in turn increased student motivation and predicted course grades [73]. Hearing student voices when answering in-class questions also has an impact on how students view the academic ability of their peers. A study by Grunspan et al., found that males underestimated the academic ability of their female classmates in courses where males were more outspoken than females during class [74]. To ensure that all student voices are heard, instructors can use random or warm calling when calling on students [75,76].

We were surprised that there was no evidence to show a positive or negative correlation for the use of high intensity of the learner-centered practices with exam scores. One possible explanation for this result could be that in our sample, there were more upper-division courses in the fourth quartile than in the other three quartiles. To enroll in upper-division science courses at this university students must successfully advance through the introductory level prerequisite courses. It is possible that students in these upper-division courses have become more autonomous learners and are not as dependent on the teaching methods used in the courses.

#### Introductory STEM courses

Students in introductory STEM courses are usually earlier in their academic careers and may require more scaffolding and support in the course. Indeed, we found that students in these courses also benefited from the medium intensity of learner-centered practices as their exam scores also increased. This result could indicate that these practices provide the level of support younger students need to perform well on exams. Interestingly, students in introductory courses did not benefit from the medium-high or high levels of learner-centered practices. It’s possible that these students didn’t have enough prior experience with these new teaching methods, which may have prevented them from fully benefiting from their impact on learning. These results could be encouraging to faculty of introductory courses who are just learning how to implement learner-centered teaching practices. Our results indicate that even a medium level of implementation can benefit students.

### RQ2a. Do the correlations vary by student demographic groups?

#### Gender

Another aspect of second-generation active learning research is to explore the impact of the teaching intervention on the disaggregated student population. Many studies have reported gender gaps in academic performance in STEM courses [71,75,77–80]. We were surprised to see a gender difference at medium levels of implementing learner-centered teaching practices. Our results indicate that male students’ exam performance benefited more than female students’ in courses taught at a medium level. In more recent years, there has been a growing concern that boys and young men are being left behind by the educational system as fewer males are graduating from high school and fewer males are earning college degrees as compared to females [81]. Men may be entering college more underprepared for learning than their female colleagues. Learner-centered teaching practices are designed to scaffold and model the learning process for students. Therefore, underprepared students may benefit most from these teaching practices in small doses.

We were encouraged to see that there was no significant difference between exam performance of male and female students across courses taught at a medium-high or high intensity of learner-centered practices compared to low levels of learner-centered practices. Therefore, we can conclude that learner-centered teaching practices are equally effective at enhancing exam performance for both groups of students at these two levels of use.

#### First Generation Status

Numerous studies have documented the academic gap that many now refer to as an opportunity gap that exists for first generation students [2,82,83]. Many explanations have been offered to explain this gap, from poor preparation in high schools [84], to a decreased sense of belonging in STEM courses [83], to systemic factors of course structure in higher education [82]. To ameliorate some of these factors, many have proposed incorporating a higher intensity of active learning teaching techniques in higher education [2,64,85]. We were pleased to see that both a medium and a medium-high level of learner-centered teaching are associated with higher exam scores in both first generation and non-first generation students. Just as with gender, the learner-centered teaching methods were equally effective with these two groups of students. It was encouraging to see that a high level of learner-centered had a significantly greater correlation with exam scores for first generation students than for non-first generation students, though the exam score increase by itself is not significantly different from zero.

### Limitations

We were not able to determine the cognitive challenge of the exams used in the courses in this study given the diversity of content across the seven STEM departments and the lack of knowledge of the specific topics that were explicitly taught in each course. Most exams in STEM across US institutions are a lower Bloom’s level [86,87], which are often much easier in terms of difficulty than higher Bloom’s level questions. It is possible the higher exam scores in some courses are due to a lower cognitive challenge level of the exam, but we have no reason to believe that easier exams are more correlated with more learner-centered teaching practices.

Another possible limitation of our study is the classroom observation tool used in the study. The PORTAAL tool was developed in biology classrooms, though the practices included in PORTAAL are not limited to practices used in biology. We did however find that it was more difficult to categorize some of the teaching practices that occurred in CSE and mathematics courses because of the course content and the types of active learning practices those disciplines tend to use. We found that PORTAAL could still be adapted for these classrooms, but it is important to have conversations as a research team and come to a consensus about how to code these discipline-specific types of activities.

To investigate the association between teaching practices and exam score, ideally we would want the courses to only differ in teaching practices but be otherwise similar. However, in practice, many aspects of the courses could be different, such as the amount of homework assigned, the availability of teaching or learning assistants, problem solving sessions, and office hours. Most of these aspects would contribute to student exam performance, while being correlated with in-class teaching practices at the same time. Although the random effects included in our model could partially account for the intrinsic differences between courses, it appears that certain important course-level confounders such as exam difficulty or work completed outside of class time would still need to be adjusted in order to achieve a more comprehensive conclusion. Therefore, we need to be cautious when interpreting the significant results from our current models and acknowledge the possibility of spurious associations caused by unmeasured confounders.

## Conclusions

Our work is a significant contribution to the new field of second-generation active learning research. We have extended our previous research in this field by analyzing sets of evidence-based teaching practices (PORTAAL practices) rather than isolated practices used in the classroom. We have shown that the use of the set of practices termed “learner-centered” correlates with higher exam scores as compared to “instruction-centered” practices. Furthermore, we have evidence that as the intensity of use of learner-centered practices increases, so too does student exam performance. We have also expanded our previous findings on the positive correlation of the use of evidence-based teaching practices with student exam performance from one STEM department, biology, to six other STEM departments: chemistry, CSE, mathematics, physics, psychology, and public health and from primarily lower-division courses in biology to upper- and lower-division courses across these seven STEM disciplines. Our results support the claim that learner-centered practices improve students’ academic performance across many STEM disciplines and across academic levels.

By using the PORTAAL classroom observation tool to analyze the courses in our study, we have been able to identify at a more granular level the characteristics of teaching methods that are effective at improving exam performance. As most named active learning teaching practices require fidelity of implementation, often having multiple steps or tasks that should be completed in a specific order, this can be daunting for instructors to execute. The results from our study may support faculty as they mix and match the use of PORTAAL practices based on their own comfort level with the practices and implement practices that are appropriate for the content area of their course and the abilities of their students.

As the evidence accumulates that increasing student engagement in the learning process enhances student academic performance and narrows opportunity gaps for underrepresented students [2,64,85] it may be time to change the term “opportunity or achievement gap” to “teaching gap”, as there is now significant evidence in the literature that teaching practices have a demonstrable impact on student academic performance.

## Acknowledgements

We thank the undergraduate researchers who used PORTAAL to code the classroom recordings: E. Swanberg, P.J. Kambhiranond, A. Immel, C. Meng, and J. McAleer.

## Supporting information

**S1 Appendix. PORTAAL coding rubric.**

**S2 Appendix. PORTAAL rubric manual.**

**S3 Table. PORTAAL practice code names and long descriptions, intensity or duration.**

**S4 Table. Number of courses, instructors, and student data points for each STEM department.**

**S5 Table. The number (percentage) of lower- and upper-division courses in each quartile of PC2.**

**S6 Table. Descriptive statistics of student data points in all courses by demographic variables.**

**S7 Fig. Correlation matrix of the 10 PORTAAL practices used for analysis.**

**S8 Fig. Scree plot of the percent variance explained by each of the 10 principal components (PCs) included in the principal component analysis.**

**S9 Table. LMM results for introductory courses.**

**S10 File. Full dataset for analysis.** Includes average student exam scores, demographic information, course information, and mean PORTAAL practice values for all student data points.

